# New Measures of Visual Scanning Efficiency and Cognitive Effort

**DOI:** 10.1101/2020.11.17.386185

**Authors:** Zezhong Lv, Qing Xu, Klaus Schoeffmann, Simon Parkinson

## Abstract

Visual scanning plays an important role in sampling visual information from the surrounding environments for a lot of everyday sensorimotor tasks, such as walking and car driving. In this paper, we consider the problem of visual scanning mechanism underpinning sensorimotor tasks in *3D* dynamic environments. We exploit the use of eye tracking data as a behaviometric, for indicating the visuo-motor behavioral measures in the context of virtual driving. A new metric of visual scanning efficiency (*VSE*), which is defined as a mathematical divergence between a fixation distribution and a distribution of optical flows induced by fixations, is proposed by making use of a widely-known information theoretic tool, namely the square root of *Jensen-Shannon divergence*. Based on the proposed efficiency metric, a cognitive effort measure (*CEM*) is developed by using the concept of quantity of information. Psychophysical eye tracking studies, in virtual reality based driving, are conducted to reveal that the new metric of visual scanning efficiency can be employed very well as a proxy evaluation for driving performance. In addition, the effectiveness of the proposed cognitive effort measure is demonstrated by a strong correlation between this measure and pupil size change. These results suggest that the exploitation of eye tracking data provides an effective behaviometric for sensorimotor activity.

## I. Introduction

### A. Background

Visual scanning is critical for the living of any human in natural surroundings [1]. Visual scanning is indeed the cornerstone for a human to perform common and everyday sensorimotor tasks, such as walking and car driving. Actually, the understanding of the mechanism behind visual scanning has been very critical since late 1970 and, is especially helpful and beneficial to making stark and essential progress in both theoretical and practical perspectives [2]. As has been widely accepted, visual scanning in sensorimotor activities is a typical task directed behavior contributed by top-down modulation [3]. And this is the focused consideration in this paper.

Basically, it is significant to make clear how well a human can achieve for sampling visual information through visual scanning. In this case, to define an efficiency measure objectively is one of the most fundamental points involved in the understanding of visual scanning mechanism [2]. Also it is notably argued that the measure of efficiency for visual scanning should be indicative for the performance of the corresponding sensorimotor task [4].

But unfortunately, all the relevant research findings available do have some issues, which will be described in Section II. Indeed, it is challenging to measure the visual scanning behavior, due to that the behavior itself, which results from both the bottom-up influence by visual stimuli and the top-down modulation impact by human activity, is very sophisticated, according to the theory of predictive processing and active inference [5]. Besides, there is no established work on defining a measure of visual scanning efficiency as a true math metric, and obviously this remarkable mathematical property makes the measure more usable in theory and practice [6]. In order to combat all the disadvantages mentioned above, a metric of visual scanning efficiency is proposed based on making use of the information theoretic tools, behaving very well as a proxy for the driving performance in the context of virtual driving.

### B. Contributions

The main contributions of this paper are as follows.

- The optical flow induced by a fixation is introduced, based on the detection of the visual motion resulted from a fixated dynamic stimulus.
- We propose a methodology to define a mathematical metric of visual scanning efficiency in an objective and quantitative way, by the square root of *JSD* divergence between fixation distribution and optical flow distribution.
- Based on an extended exploitation of our proposed visual scanning efficiency, a cognitive effort measure, in a perspective of the quantity of information, is developed.

To the best of our knowledge, this is the first time to effectively define the metric of visual scanning efficiency and cognitive effort measure, based on the exploitation of information theoretic tools. This paper paves a novel path for behaviometric discovery by the utilization of eye tracking data.

### C. Organization

Firstly, the related works are discussed in Section II. The philosophy and methodology of our proposed technique are detailed in Section III. Then the experimental methods are described in Section IV, followed by the results and discussion presented in Section V. Finally, conclusions and future works are given in Section VI.

## II. Related Works

The initial consideration for discussing the efficiency measure of visual scanning is to select suitable eye tracking indices naturally associated with cognitive processing. Fixation and saccade are classic indices for this purpose. But, direct and indirect usages of these indices (for examples, rate/duration of them and their simple combinations) are more applicable in specific application scenarios [7], than in the general efficiency definition. In addition, pupil dilation and blink rate are two widely used eye tracking indices for the study of cognition and psychology [8].

The efficiency evaluation of visual scanning has started from the study of the complexity of visuo-motor behavior and, this is the usual manner in relevant field [2]. Information entropy, which is a good measure of complexity [9], in effect has been used a lot for the issue of visual scanning complexity [2]. The so-called entropy rate, originally used for the measurement of task load [10], is defined as a multiplication between the normalized entropy of fixation sequence and the sum of inverse transition durations. Entropy of fixation sequence (called *EoFS* for brevity) defines based on the probability distribution of fixation sequences, giving the degree of complexity for spatial patterns of fixation sequence [11]. *EoFS* has shown its interaction to visual content complexity [12] and to human factors such as expertise skill [11].

Differently from *EoFS*, gaze transition entropy (*GTE*), in which transition between fixations is depicted by the probability transition between *Markov* states (in this paper, meaning the areas of interest (*AOI*s)) and, conditional entropy based on transition probabilities is used for its definition to express the degree of complexity for temporal patterns of fixation transition, with respect to overall spatial fixation dispersion. *GTE* has demonstrated its relationship with visual content complexity [13] and also with human factors, such as anxiety [14]. There exists a closely relevant gaze entropy, called stationary gaze entropy (*SGE*) [13], in which the *Shannon* entropy, based on an equilibrium probability distribution for *Markov* transition matrix, is used to describe the degree of complexity for spatial patterns of fixation dispersion. *SGE* has also exhibited, similar to *GTE*, its connection to visual content complexity [13], and also to human factors, such as emotional valence [15].

Notice specifically that, it is necessary to know the relationship between entropy based measures discussed above and the performance of sensorimotor tasks. *EoFS* has not been indicated to have correlation with the task performance. It has been reported that *GTE* [1] could not predict the performance of sensorimotor tasks. As for *SGE*, it relates to the complexity of visual content, but can not indicate the performance of task directed sensorimotor activities [3]. As a mater of fact, eye tracking has been expected as a strong estimator for task performance in many professional fields [4]. Obviously, it is indeed indispensable for the efficiency measure of visual scanning to take the role as measure of task performance.

## III. METHODOLOGY

### A. The Philosophy of the Proposed VSE *and* CEM

It is well understood that, in a sensorimotor task, visual scanning is managed by a performer through a series of fixations, so that meeting the requirement of visual sampling the surrounding environments for fulfilling this task [16]. The probability distribution of fixations, namely the fixation distribution, which is constructed based on the normalized histogram of fixation locations in a *3D* environment, is used as a basis for understanding the mechanism of visual scanning in this paper. In addition, the optical flow, which effectively reflects a visual motion resulted from a dynamic stimulus [17], always happens in driving situations [18]. In this paper, the optical flows induced by fixations are detected and used to establish the optical flow probability distribution. And then a divergence between fixation distribution and optical flow distribution is proposed to define the visual scanning efficiency. Notice that, optical flow represents a vector, according to its original definition [17] [18]; but, for the purpose of this paper, unless otherwise specified, optical flow is always used for representing its magnitude.

The rationale behind our proposed divergence comes from two points. Firstly, optical flow distribution in fact contains the visual motions that are embedded in fixation distribution, the divergence between these two distributions represents visual motions perceived by a performer in visual scanning. Secondly, bear in mind that visuo-motor behavior is usually influenced by both top-down and bottom-up factors and that, the combination of these two factors reflects the visual scanning efficiency [2]. In fact, all the visual motions, which are essential information due to the dynamic nature of driving environment, are considered to result from bottom-up factors [2]. To go further, visual motions perceived by a performer in the procedure of driving actually indicate an effective component of all the visual motions necessarily used for the purpose of fulfilling the driving task. As a result, this so-called effective component can be easily considered to be caused by both top-down modulations and bottom-up influences. And in this case, we hypothesize that the proposed divergence takes as an indicator for visual scanning efficiency.

The square root of *JSD* (*SRJSD*) is chosen for computing the divergence and for extracting the visual motions, because it is largely accepted as a true mathematical metric [19]. Actually, *SRJSD* offers a kind of distance between two probability distributions in a normalized manner [6] [19]. Thus, *SRJSD* between fixation distribution and optical flow distribution in effect indicates a normalized degree to which the visual motions are perceived. To sum up, *SRJSD* between fixation and optical flow distributions should behave as an evaluation function for visual scanning efficiency (*VSE*). The larger the *SRJSD*, the more visual motions are perceived, the higher the *VSE* becomes, and *vice versa*.

Additionally, the proposed *VSE* can be studied from the perspective of probability and information theories. That is, the *VSE*, in fact, can be considered as a probability *p* an event occurs, because it ranges from 0 to 1. As a result, the *VSE* points out the probability of perception of the visual motions. According to information theory, the logarithmic probability of occurrence (− log_2_ *p*) represents the quantity of information conveyed by the occurrence [9]. We hypothesize that the information quantity may indicate the attentional resources needed for the perception of all the visual motions. Because the amount of attentional resource reflects cognitive effort [8], and thus the logarithm of *VSE* may be the central part of cognitive effort.

A scheme of the calculation of the two proposed measures is shown in Fig. 1. Firstly, optical flows induced by fixations are extracted, to construct the optical flow distribution that actually embeds visual motions into the fixation distribution. Then the *VSE* is obtained by *SRJSD* between these two distributions, for measuring the perceived visual motions in driving. Finally, the *CEM* is obtained by the logarithm of *VSE*.

**Fig. 1.**
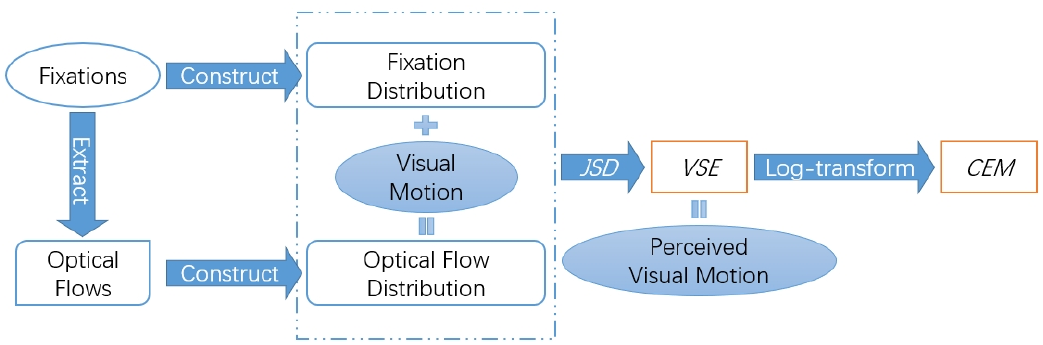
The scheme of the calculation of the proposed measures.

### B. Optical Flow Induced by Fixation

A fixation *f*_*n*_ (its index is *n*) with duration *τ*_*n*_ by an observer *O* in a *3D* environment is shown in Fig. 2; in this figure, *ρ*_*n*_ is the distance between *O* and *f*_*n*_ in the direction of the line of sight, and *ρ*_*n*_ is the depth of *f*_*n*_ from the perspective of O. Considering there is a relative displacement ***u***_***n***_, during *τ*_*n*_, between *O* and *f*_*n*_, ***υ***_***n***_ is defined as the optical flow vector [17] (bold-type letters are used to denote vectors in this paper). Actually, ***υ***_***n***_ is the projection of ***u***_***n***_ in the direction of optical flow vector, and thus perpendicular to the line of sight. The so-called visual motion perceived by the observer, *m*_*n*_, is defined as the trajectory segment of the circular motion of *f*_*n*_ centered on *O* in this paper. Note, *m*_*n*_ acts as an approximated magnitude of ***υ***_***n***_. The central angle *I*_*n*_ subtended by *m*_*n*_,

**Fig. 2.**
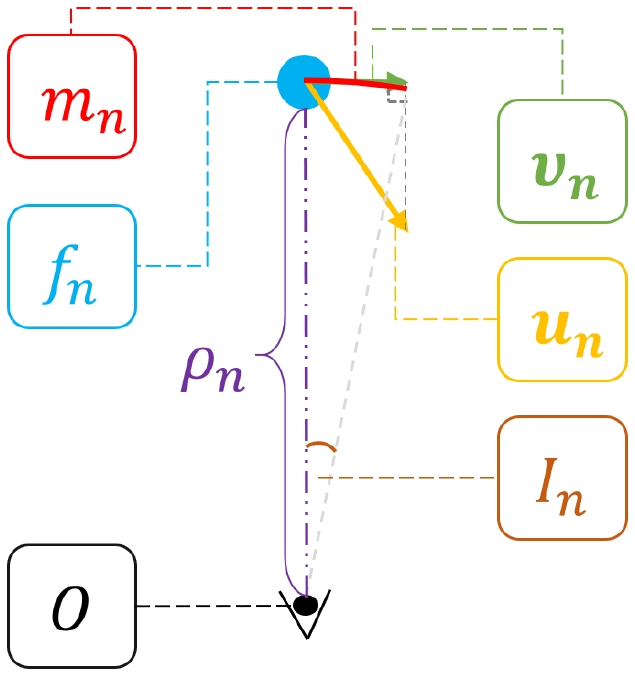
An illustration of optical flow *I*_*n*_ induced by fixation *f*_*n*_.

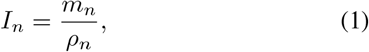

is used to approximately define a perceived magnitude of optical flow, according to the usual practice that angle is widely used to represent the magnitude in eye tracking [20]. Notice that, the definition of the perceived magnitude of optical flow explicitly uses the depth, enabling it to convey an important information in *3D* environments. For the sake of this paper, *I*_*n*_ is called as optical flow.

### C. Visual Scanning Efficiency

In this paper, fixations are assigned to *AOI*s, and *AOI*s are defined based on *3D* objects in virtual environment [21]. There exists a one-one correspondence between the *AOI*s and the objects.

The probability distribution of fixations, which is the normalized fixation histogram, is based on the frequencies of fixations located at *AOI*s. There exist a series of fixations *F* = ⟨*f*_1_, …, *f*_*n*_, …, *fN*_*F*_⟩, (*n* ∈ *C*_*F*_ = {1, …, *N*_*F*_}) that happens in a driving procedure, the *AOI*s hit by these fixations are *X*_*F*_ ={*χ*_*i*_| *i* = 1, …, *N*_*A*_}. For an *AOI χ*_*i*_ ∈ *X*_*F*_, all the indices of the fixations {*f*_*h*_ |*h* ∈ *C*_*F*_} hitting *χ*_*i*_ constitute a set,

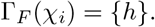

Then the fixation distribution *P*_*f*_ = {*p*_*f*_ (*χ*_*i*_) | *χ*_*i*_ ∈ *X*_*F*_} and the optical flow distribution *P*_*o*_ = {*p*_*o*_(*χ*_*i*_) | *χ*_*i*_ ∈ *X*_*F*_} are obtained respectively as

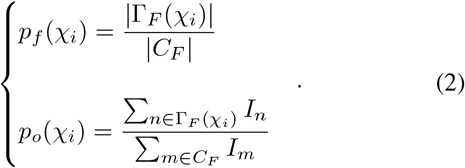

It is noted that, visual motions, indicated by optical flows, are embedded into the fixation distribution *P*_*f*_ to construct the optical flow distribution *P*_*o*_.

A special instance of *JSD* between two distributions {*P* = *p*(*x*) | *x* ∈ *X* and *Q* = {*q*(*x*) | *x ∈ X*} with equal weights [6] is defined as

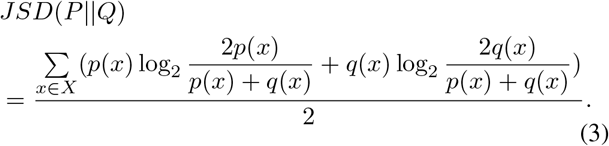

The square root of *JSD*(*P* || *Q*)

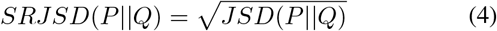

is used to obtain the metric of visual scanning efficiency,

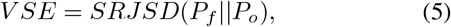

because *SRJSD* is a true mathematical metric [19].

### D. Cognitive Effort Measure

Because the distributions of fixation and fixation sequence reflect different aspects of visual scanning, a combination of both these two distributions should provide a more complete understanding of visual scanning behavior, as has been pointed out in relevant work [2]. Therefore, we propose to utilize the fixation sequence distribution for the sake of a scaling function, detailed in the following text of this section.

There exist a series of fixations *F*_*S*_ = ⟨*f*_1_, …, *f*_*n*_, …, *fN*_*f* +1_ ⟩, then we have *N*_*f*_ fixation sequences *S* = {*f*_⟨*n,n*+1⟩_ = ⟨*f*_*n*_, *f*_*n*+1_⟩ | ⟨*n, n* + 1⟩ ∈ *C*_*S*_}, *C*_*S*_ = {⟨1, 2⟩ …, ⟨*N*_*F*_, *N*_*F*_ + 1⟩}. A set of *AOI* sequences is constituted as *X*_*S*_ = {*χ*_⟨*i,j*⟩_ = ⟨*χ*_*i*_, *χ*_*j*_ ⟩ | ⟨*i, j*⟩ ∈ *A*_*S*_}, where *A*_*S*_ contains the indices of all the *AOI* sequences *χ*_⟨*i,j*⟩_ hit by all the fixation sequences in *S*, note that a fixation sequence *f*_⟨*n,n*+1⟩_ hitting *AOI* sequence *χ*_⟨*i,j*⟩_ indicates a transition from *f*_*n*_ on *AOI χ*_*i*_ to *f*_*n*+1_ on *AOI χ*_*j*_. This means that the performer of visual scanning voluntarily exerts some cognitive efforts to switch the fixations from *f*_*n*_ to *f*_*n*+1_. And in this case, the optical flow *I*_*n*_ induced by *f*_*n*_, which is taken for representing the cognitive efforts exerted, is considered lost for the purpose of completing the transition of fixations from *f*_*n*_ to *f*_*n*+1_. So in this paper, for each *AOI* sequence *χ*_⟨*i,j*⟩_ ∈ *X*_*S*_, all its corresponding optical flows induced by *f*_⟨*n,n*+1⟩_ satisfying that *f*_⟨*n,n*+1⟩_ hits *χ*_⟨*i,j*⟩_ are obtained, here each optical flow takes *I*_*n*_ and is denoted as *I*_⟨*n,n*+1⟩_. Then for all the *AOI* sequences in *X*_*S*_, their corresponding optical flows are embedded into the fixation sequence distribution to construct the corresponding optical flow distribution. Note, the *SRJSD* between these two distributions takes as the scaling function for defining the cognitive effort measure.

For an *AOI* sequence *χ*_⟨*i,j*⟩_ ∈ *X*_*S*_, all the indices of the fixation sequences *f*_⟨*l,l*+1⟩_ ∈ *S* hitting *χ*_⟨*i,j*⟩_ constitute a set

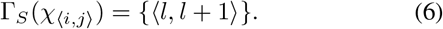

Then the distribution of fixation sequences *P*_*fs*_ = {*p*_*fs*_(*χ*_⟨*i,j*⟩_)|*χ*_⟨*i,j*⟩_ ∈ *X*_*S*_} and its corresponding optical flow distribution *P*_*os*_ = {*p*_*os*_(*χ*_⟨*i,j*⟩_|*χ*_⟨*i,j*⟩_ ∈ *X*_*S*_} are obtained as

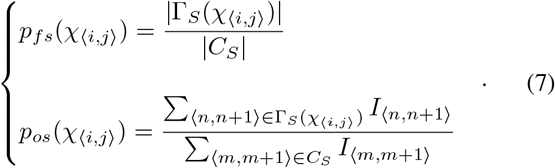

It is worth pointing out that if an *AOI* sequence *χ*_⟨*i,j*⟩_ is not hit by any fixation sequence, then this sequence will not be included in *X*_*S*_. Therefore, it is easy to see that

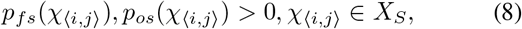

this means that the probability mass functions of fixation sequence distribution and its corresponding optical flow distribution are strictly positive.

For a certain *AOI χ*_*i*_∈ *X*_*F*_, we have the quantitative relationships that

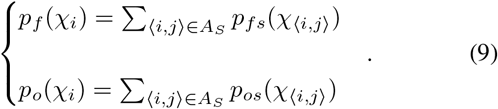

Notice that *SRJSD* between two probability distributions *P* and *Q* is not smaller than that between the distributions constructed by the sums of the corresponding subsets of *P* and *Q* respectively, as proved in Theorem 1.

#### Theorem 1

There are two probability distributions *P* = {*p*(*x*)|*x* ∈ Ψ} and *Q* = {*q*(*x*)|*x* ∈ Ψ} containing non-empty and non-overlapping subsets *µ*_*j*_ and *ν*_*j*_ (*j* = 1, …, *M*) respectively. Here *µ*_*j*_ = {*p*(*x*)|*x* ∈ *ψ*_*j*_, *ψ*_*j*_ ⊆ Ψ} and *ν*_*j*_ = {*q*(*x*)|*x* ∈ *ψ*_*j*_, *ψ*_*j*_ ⊆ Ψ}. Then two sets can be constituted as 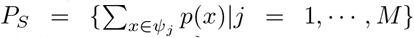 and 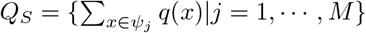. Then we have

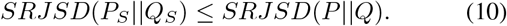

**Proof**. The log sum inequality [9] says that, for nonnegative numbers *a*_1_, …, *a*_*n*_ and *b*_1_, …, *b*_*n*_,

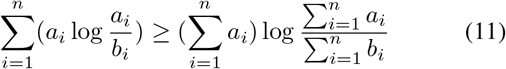

holds, here equality happens if and only if 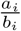 is a constant.

Setting *a* = *p*(*x*), *x* ∈ *ψ*_*j*_ and 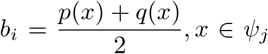, we obtain

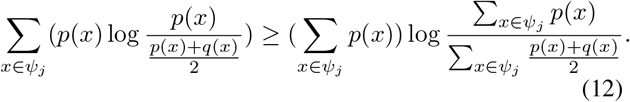

Similarly,

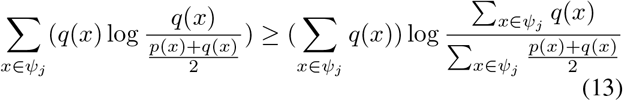

can be obtained. Thus,

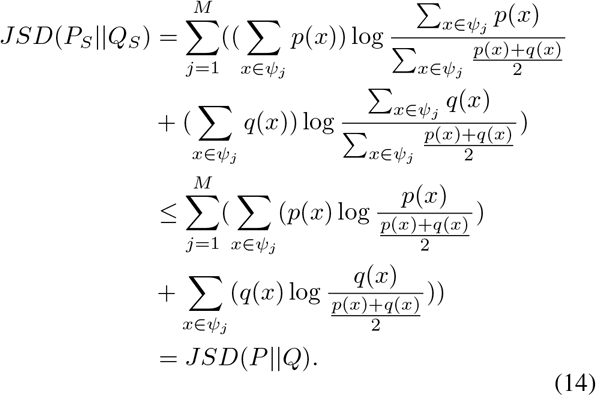

Therefore we have

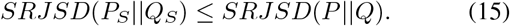

When the fixation distribution and the optical flow distribution, as shown in (9), are applied to Theorem 1, we obtain

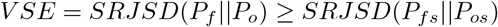

and

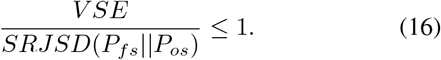

Therefore, *SRJSD*(*P*_*fs*_ ||*P*_*os*_) is considered as a normalization factor for *VSE* in this paper.

As discussed in Section III-A, the central part of *CEM* is −log_2_ *V SE*. Inspired by (16), we propose the hypothesis that the cognitive effort measure (*CEM*) takes a scaled form

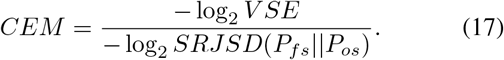

We set *CEM*= 0 when *JSD*(*P*_*fs*_ ||*P*_*os*_) = 0.

Notice that *JSD* between two probability distributions with strictly positive probability mass functions is always lower than 1, and this is proved in Theorem 2.

#### Theorem 2

There are two probability distributions *P* = {*p*(*x*) |*x* ∈ Ψ} and *Q* = {*q*(*x*) | *x* ∈ Ψ}, where *p*(*x*), *q*(*x*) *>* 0, then we have *JSD*(*P*||*Q*) *<* 1.

**Proof**.

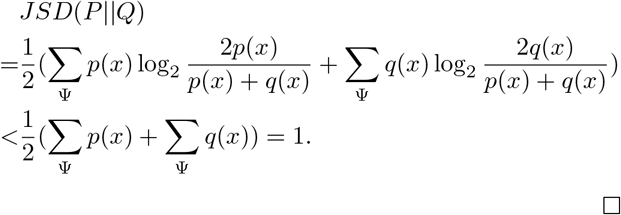

Because of the positive property of the two distributions *P*_*fs*_ and *P*_*os*_ given by (8), according to Theorem 2, we have log_2_ *SRJSD*(*P*_*fs*_ ||*P*_*os*_) ≠ 0. And thus, *CEM* always makes sense.

### IV. EXPERIMENT

#### A. Participants

Fourteen Master/Phd students (5 females; age range: 21-29, Mean = 21.3, SD = 2.37) with driving experience (they hold their driver license at least one and a half years) from Tianjin University volunteer to participate in the psychophysical studies. All of the participants have normal/corrected-to-normal visual acuity and normal color vision. There is no participant having adverse reaction to the virtual environment we set up for the studies.

### B. Virtual Reality Environment

In order to deeply understand visuo-motor behavior based on eye movement and driving data in an effective and controllable way, virtual reality is used in this paper, as has been done in many works [22]. A virtual environment (*VE*), including common straight, curved roads and buildings, is designed for the experimental study. In this paper, visual scanning happened for sensorimotor tasks is considered as a task directed behavior, as has been widely done in many relevant works [16] [3]. Therefore, there are no visual stimuli based distractors, such as pedestrians, traffic light signals and similar, are included in the *VE*.

### C. Apparatus

*HTC Vive* headset [23] is used to display the *VE* for participants. The eye-tracking equipment is *7INVENSUN Instrument aGlass DKII* [24], which is embedded into the *HTC Vive* display to capture visual scanning data in a frequency of 90 Hz and in an accuracy of gaze position of 0.5?. The driving device is a *Logitech G29* steering wheel [25]. Participants listen the ambient traffic and car engine sounds in *VE* by speakers. The visual scanning and driving behaviors of participants are displayed on desktop monitor to observe the experimental procedure.

### D. Driving Task

Driving is taken as the sensorimotor task, because it is typically happened in everyday life. As discussed in Section IV-B and, as well as in relevant research [16] [3], the task directed focus on visual scanning and driving is taken. And as a result, participants are required to keep driving at a target speed of 40 km/h, for speed control. This paper takes the inverse of the mean acceleration of vehicle to denote the driving performance. The smaller the mean acceleration, the higher the driving performance becomes, and *vice versa*. In fact, this kind of performance measure has been used a lot in literature [26]. An example of performing driving tasks is presented in Fig. 3.

**Fig. 3.**
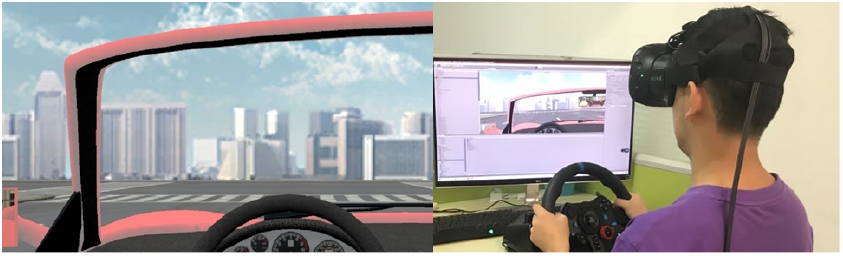
A participant is performing the driving task in the *VE*.

### E. Procedure

Each participant completes four test sessions with the same task requirements and the same driving routes, with an interval of one week between every two sessions. All the sessions for each participant begin with a 9-point calibration for the eye tracker. In this paper, a trial represents a test session, and there are 14*4 = 56 valid trials in all. Data for visual scanning and driving behaviors are recorded during test sessions.

A preparation session is applied to participants before each test session to let them know the purpose and procedure about the psychological studies.

## V. Results and Discussions

### A. Correlation between VSE and Driving Performance

We verify the hypothesis that the proposed *VSE* takes as an indicator for visual scanning efficiency. Because visual scanning efficiency is usually expected to reflect the performance of sensorimotor tasks [2], correlation analysis based on three kinds of widely used correlation coefficients (*CC*) [27], namely Pearson linear correlation coefficient (*PLCC*), Spearman rank order correlation coefficient (*SROCC*) and Kendall rank order correlation coefficient (*KROCC*), is employed to validate the proposed hypothesis. As listed in Table I, the *CC* results show a statistically significant correlation between our proposed efficiency measure and the task performance. But in contrast, the entropy based measures and eye tracking indices do not correlate with the driving performance. These findings suggest that the new proposed efficiency metric may be a proxy assessment for the driving performance in virtual environment. An example of eye tracking and driving performance data for two trials, Trials A and B, from the second and fourth test sessions conducted by the 4th and 13th participants, respectively, are presented in Fig. 4. This figure has upper and bottom components, the probability distribution histograms and the fixation-acceleration plot. In the probability distribution histograms of Trials A and B, blue and yellow bars indicate the fixation distributions and optical flow distributions, respectively. Almost all of the differences between the correspondence blue and yellow bars in Trial A are larger than those in Trial B; in particular, for example, this happens for *AOI*s 2 and 5. Note *JSD* is calculated as a sum of the differences between the correspondence probabilities of two distributions. Obviously, in this case, the divergence between the fixation and optical flow distributions from Trial A (*SRJSD*=0.23) is largely bigger than that from Trial B (*SRJSD*=0.01).

**TABLE I.**
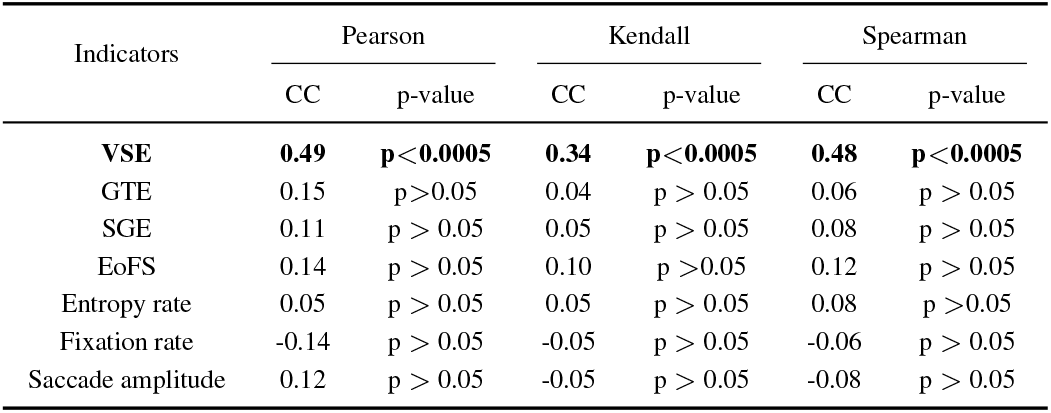
CC between indicators and driving performance

| Indicators | Pearson |  | Kendall |  | Spearman |  |
| --- | --- | --- | --- | --- | --- | --- |
|  | CC | p-value | CC | p-value | CC | p-value |
| VSE | **0.49** | **p<0.0005** | **0.34** | **p<0.0005** | **0.48** | **p<0.0005** |
| GTE | 0.15 | p>0.05 | 0.04 | p>0.05 | 0.06 | p>0.05 |
| SGE | 0.11 | p>0.05 | 0.05 | p>0.05 | 0.08 | p>0.05 |
| EoFS | 0.14 | p>0.05 | 0.10 | p>0.05 | 0.12 | p>0.05 |
| Entropy rate | 0.05 | p>0.05 | 0.05 | p>0.05 | 0.08 | p>0.05 |
| Fixation rate | -0.14 | p>0.05 | -0.05 | p>0.05 | -0.06 | p>0.05 |
| Saccade amplitude | 0.12 | p>0.05 | -0.05 | p>0.05 | -0.08 | p>0.05 |

**Fig. 4.**
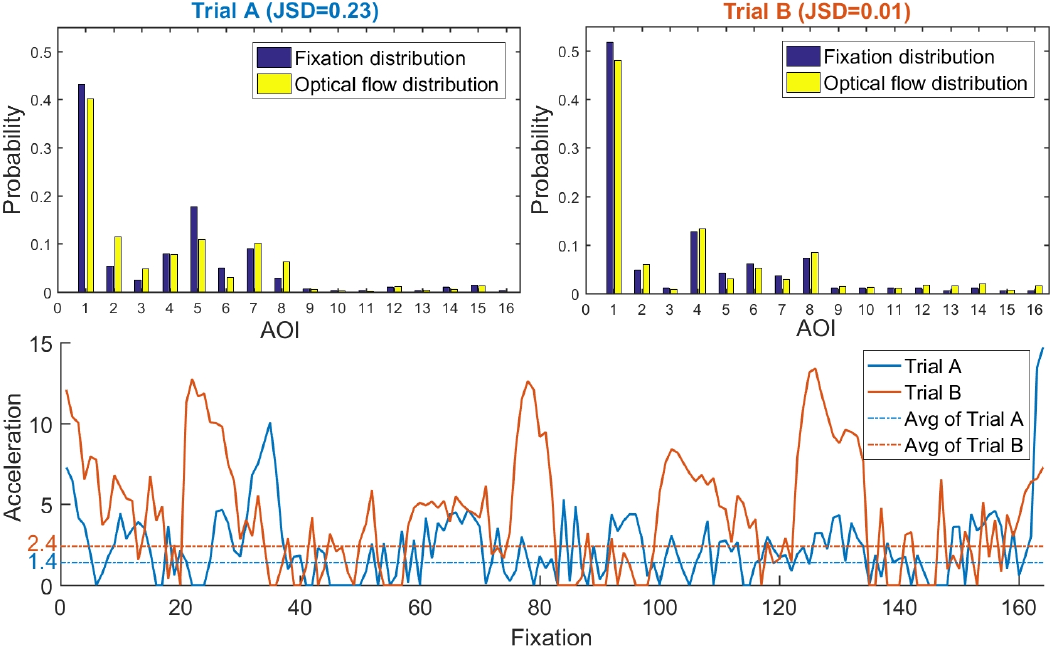
Illustrations of the fixation and optical flow distributions from Trials A and B, *SRJSD* = 0.23 of Trial A is much higher than *SRJSD* = 0.01 of Trial B (upper). The driving performance of Trial A is largely better than that of Trial B, as shown by their average mean accelerations 1.4 and 2.4, respectively (bottom). The *JSD* visual scanning efficiency corresponds very well to the driving performance.

In the fixation-acceleration plot, the horizontal and vertical axes represent the indices of fixations and the accelerations, respectively. Average accelerations of Trials A and B are 1.4 and 2.4, which are indicated by the blue and orange dashed lines, respectively. Therefore, task performance of Trials A is much better than that of Trial B. This means that, a higher *SRJSD* corresponds well to a lower acceleration and thus, to a better driving performance, and *vice versa*, validating the effectiveness of the proposed metric of visual scanning efficiency.

In addition, a linear regression analysis is performed for the further confirmation of the correlation, as shown in Fig. 5, where the horizontal and vertical axes represent the proposed *VSE* and the driving performance (inverse of mean acceleration) respectively. The regression result shows a significant linear relationship between the proposed efficiency metric and driving performance.

**Fig. 5.**
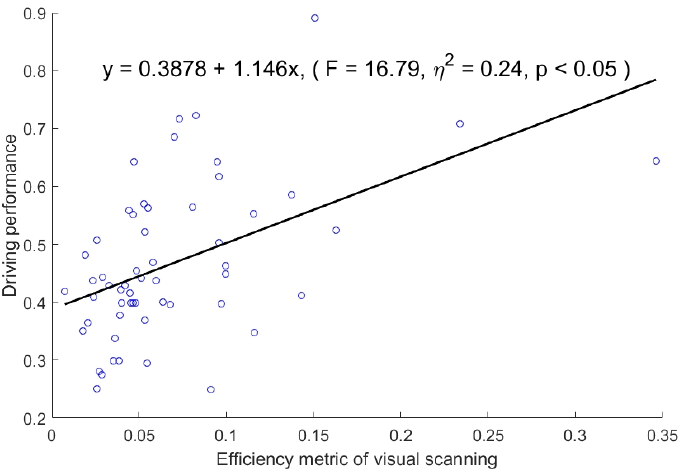
Linear regression analysis between the *VSE* and driving performance.

### B. Correlation between CEM and Pupil Size Change

Pupil size change, as a kind of pupillary response, is an autonomic and reflexive indicator of cognitive effort [8] [28]. In this paper, we utilize the standard deviation of pupil size during each trial to represent the pupil size change because of its simplicity and effectiveness [29]. A correlation is revealed between the cognitive effort measure and the pupil size change, as listed in Table II, which validates the hypothesis that the proposed *CEM* can be an indicator for cognitive effort.

**TABLE II.**
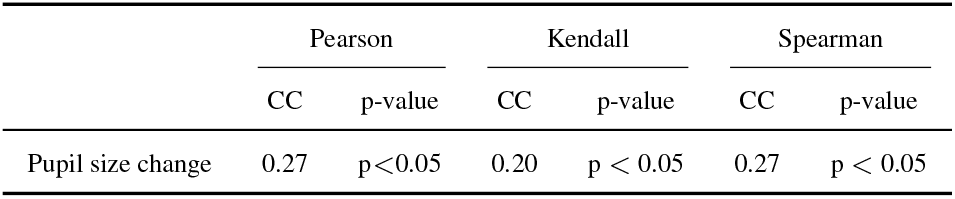
CC between CEM and pupil size change

|  | Pearson |  | Kendall |  | Spearman |  |
| --- | --- | --- | --- | --- | --- | --- |
|  | CC | p-value | CC | p-value | CC | p-value |
| Pupil size change | 0.27 | $p < 0.05$ | 0.20 | $p < 0.05$ | 0.27 | $p < 0.05$ |

### C. Discussions

This paper discovers a behaviometric based on the exploitation of eye tracking data, for investigating and understanding the efficiency of visual scanning and the cognitive effort, in the procedure of driving. To examine these findings, participants are asked to drive at a target speed in a *3D* virtual environment. The psychophysical results have clearly suggested that the proposed visual scanning efficiency (*VSE*) and cognitive effort measure (*CEM*) can take good proxies for driving task performance and for pupil size change, respectively.

The *PLCC, KROCC* and *SROCC* between the proposed *VSE* and driving performance are 0.49, 0.34 and 0.48 respectively, with significant *p-value*s (*p <* 0.0005), as listed in Table I. The success of our proposed metric of visual scanning efficiency benefits from the characterization of how many optical flows (visual motions) are perceived in driving. The central consideration of this is to place the most importance on the perceived visual motions employed for the completion of visual scanning and of driving. That is, visual motion, which always and naturally happens in driving environments [18], is taken as a motivation force for visuo-motor behavior and for driving activity. And, the visual motion perceived for visual scanning during the task of driving indicates the top-down modulation exerted by driver to fulfill the task. As a matter of fact, our idea resides in that the visual scanning is a behavior directed by the driving task, as a combination of top-down and bottom-up influences; this has been widely accepted in literature [3]. As discussed above and, as commonly known in relevant research [1], the visual scanning behavior plays a very important role in performing a sensorimotor task (such as driving), this paper calculates how much correlation there exists between visual scanning efficiency and driving performance, as an evaluation of the proposed *VSE*.

The entropy based measures under comparison do not have significant correlation with task performance, and the direct use of eye tracking indices (fixation rate and saccade amplitude) does not either. Indeed, the entropy based measures express the concept of complexity, which is essentially different from our proposed *VSE*. Concretely, *GTE* is a powerful estimator of the complexity for transition pattern [1] [2], but not of the efficiency for the whole visual scanning procedure. Similarly, *SGE* [13] and *EoFS* [12] give the complexity for spatial patterns of fixations and of fixation sequences, respectively. We believe that the complexity of visuo-motor behavior just plays a marginal role for the efficiency measure of visual scanning in task directed sensorimotor tasks. In fact, visual motion, which is critical for performing sensorimotor tasks like driving, is not explicitly considered by this complexity measurement. And thus this is the reason why the entropy based measures under comparison cannot relate enough to the driving task performance. The direct eye tracking indices, which are generated based on fundamental fixation and saccade, cannot be a proxy for the driving task performance. This is because that these indices are basically simple statistics and only applicable for specific applications, so as not to behave generally and robustly as an estimator of visual scanning efficiency [7].

A significant correlation (*p <* 0.05) is revealed between the proposed *CEM* and the pupil size change by the three correlation analyses, having the *PLCC, KROCC* and *SROCC* of 0.27, 0.20, 0.27 respectively, as shown in Table II. This finding verifies the power of the cognitive effort measure, according to the close association between cognitive effort and pupil size change [8] [28]. Considering that we have made progress on the exploitation of eye tracking data, as a behaviometric, for the measurements of visual scanning efficiency and cognitive effort, a further investigation into the relationship between these two proposed measures will be performed in future work. And actually, this could be a working path illuminated based on the exploitation of Yerkes-Dodson law [30].

In this paper, visual scanning in sensorimotor tasks is mainly considered as a task directed behavior, as has been largely accepted in literature [16] [3]. And in this case, two corresponding treatments are performed in our psychophysical studies. Firstly, we especially design the virtual environment not to include visual stimuli based distractors (such as pedestrians, traffic light signals and similar). The purpose of this is to use the visual motion, which is resulted from the dynamic nature of driving environment, as the single bottom-up contribution. Secondly, we simply ask participants to travel at a fixed speed as the requirement of driving task. The objective of this is to take the speed control as the single contributor of top-down modulation. As a result of these two treatments, mean acceleration is taken as the driving task performance. Note that mean acceleration is a common measure of driving performance in related research [26]. Obviously, all in all, we emphasize that the proposed *VSE* and *CEM* are investigated and assessed more easily and more clearly in the current two treatments. Complex virtual environment and more requirements of the driving task will be our future work.

Notice that the findings of this paper may not be applicable for all cases, but it does work in the context of our topic. Due to that visual scanning is exceptional important in virtual and real-world sensorimotor tasks, what we have achieved on efficiency metric and on cognitive effort measure in virtual driving should be potentially helpful for ergonomic evaluation pragmatically, in many practical and relevant applications.

## VI. Conclusions and Future Works

In this paper, we take an important step for thorough understanding the mechanism of visual scanning in virtual driving. This paper has established, in an objective and quantitative way, a new metric for visual scanning efficiency through a methodology that exploits the perceived visual motions in a sensorimotor task by the divergence between fixation distribution and optical flow distribution. The efficiency measure provides a window into how to evaluate the performance of visual scanning, and also behaves as a strong proxy for objectively assessing the performance of driving tasks. As far as we can know, no research up to now has reported this kind of finding to shed light on the issue of efficiency metric for visual scanning and for performing sensorimotor tasks. Additionally, the proposed cognitive effort measure, which is significantly correlates with pupil size change, may offer a new perspective on the inherent relationship between task directed visual scanning and eye tracking data. The two measures we have proposed have been verified by the psychophysical studies conducted in this paper. In summary, this paper proposes a new methodology to measure visuo-motor behavior by using eye tracking data, so as to help the development of behaviometric discovery, from both theoretical and practical perspectives.

In the near future, we will investigate our proposed methodology and measures for real-life driving scenarios, for instance, for crash risk problem [31]. In consideration of the critical role of illumination conditions for driving, we will exploit the manipulation of illumination levels in a detailed quantitative way, to comprehensively understand the mechanism of visual scanning efficiency. The association between the two proposed measures will be studied, by the exploitation of Yerkes-Dodson law [30], as has been mentioned in Section V-C. Also, physiological signals such as heartbeat [32] will be investigated for understanding the relationships and interplays between these signals and eye tracking data, for the sake of visual scanning efficiency and cognitive effort.

## Notes

### Competing Interest Statement

The authors have declared no competing interest.

### Summary of Updates

some typos

